# Entry Pathway for the Inverse Agonist Ligand in the G Protein-Coupled Receptor Rhodopsin

**DOI:** 10.1101/2021.05.24.445474

**Authors:** He Tian, Kathryn M. Gunnison, Manija A. Kazmi, Thomas P. Sakmar, Thomas Huber

## Abstract

While the number of high-resolution structures of ligand-bound G protein-coupled receptors (GPCRs) has been steadily climbing, ligand binding and unbinding pathways remain largely undefined. The visual photoreceptor rhodopsin (Rho) represents a curious case among GPCRs because its primary ligand 11-*cis*-retinal (11CR) is an inverse agonist, which partitions into the bilayer and is likely to enter its orthosteric binding pocket through an intermembranous pathway. Light activates Rho by converting 11CR to all-*trans*-retinal (ATR), which serves as an agonist ligand. The light-triggered switch from the inactive to the active conformation creates two openings in the transmembrane region, suggesting pathways for exit of ATR and subsequent entry of 11CR to regenerate Rho. However, stabilization of an active ligand-free opsin conformation has been found to inhibit 11CR binding. Here we address this paradox of opsin regeneration with 11CR. We used genetic code expansion to engineer Rho mutants that serve as fluorescence sensors for measuring 11CR binding kinetics and energetics. We found mutations that alter a channel between transmembrane helices 5 and 6 (TM5/6) dramatically affect 11CR binding kinetics, but not ATR release kinetics. Our data provide direct experimental evidence for 11CR entry between TM5/6 in Rho that involves dynamic allosteric control of the ligand entry channel. Our findings can be extended to other visual pigments and a wide range of GPCRs with hydrophobic ligands that are hypothesized to enter their binding pockets through transmembrane pores.

## INTRODUCTION

The visual photoreceptor rhodopsin (Rho) is G protein-coupled receptor (GPCR) that consists of an apoprotein opsin and its native ligand 11-*cis*-retinal (11CR). 11CR is covalently linked to opsin via a protonated Schiff base (PSB) bond (Wald, 1968). 11CR is a potent inverse agonist that stabilizes Rho in its inactive state with extremely low basal activity. Upon photoactivation, 11CR absorbs a photon to isomerize to all-*trans*-retinal (ATR), triggering a series of conformational changes to produce the active Meta-II Rho state capable of activating transducin. The Schiff base linkage then hydrolyzes, allowing ATR to dissociate from the ligand-binding site. In process known as “regeneration,” 11CR enters the empty ligand-binding channel of opsin to form a transient opsin-11CR complex. The formation of a PSB completes the regeneration of Rho.

Many high-resolution structures for GPCRs that bind diffusible ligands show openings connecting the extracellular surface to their orthosteric ligand-binding site. However, the crystal structures of opsin do not reveal a direct channel from the extracellular environment to the 11CR binding site. Instead, high-resolution structures of Meta-II opsin (Choe et al., 2011; Standfuss et al., 2011) and ligand-free opsin (Park et al., 2008; Scheerer et al., 2008) reveal openings in the transmembrane (TM) helical bundle at both ends of the ligand-binding pocket specific to the active conformation, suggesting that ATR exits and 11CR enters the pocket using these two openings in the TM region. Moreover, the high partition coefficient of retinal into the lipid bilayer indicates that retinal predominately exists in the lipid-solubilized form (Frederiksen et al., 2012). However, this “active conformation” model is incongruent with certain experimental observations (Schafer et al., 2015). On the other hand, recent structures of opsin bound with non-retinal small molecules reveal a channel open towards the extracellular side when the receptor was stabilized by a bulky moiety protruding between the extracellular end of TM5 and TM6 (Mattle et al., 2018). Nonetheless, it is unclear whether 11CR can induce the formation of a similar channel open towards the extracellular surface.

Here, we employ an engineered opsin that serves as a sensitive fluorescence sensor (Tian et al., 2014) for 11CR binding kinetics to identify the 11CR entry pathway. We examine how mutations designed to alter the ligand-binding channel affect the 11CR entry rate and ATR exit rate. Mutations in the channel nearest to the β-ionone end of 11CR (TM5/6) caused a ~100-fold variation in the 11CR entry kinetics, but only a 3-fold variation in the ATR release kinetics. We combined one of three TM5 mutations with I189P, which is situated in the middle of the channel and defines a key difference between rod and cone photoreceptors. The energetic additivity of the double mutant suggests that the mutations in the TM5/6 channel exert localized effects on 11CR entry. We contend that our data, and several other lines of evidence, fit best with a previously proposed model (Schafer and Farrens, 2015) in which the opsin conformer permissive for 11CR binding (Ops**) resembles that of the inactive state and the ligand binding channel needs to dynamically adjust to the incoming ligand. Our data provide direct experimental evidence for 11CR entry through a pore between TM5/6 and suggest that an “allosteric” mechanism in ligand binding in other GPCRs as well.

## RESULTS

Based on the dark-state Rho and Meta-II structures (Figure 1), we hypothesized that the residues close to the β-ionone ring end of bound retinal might play a role in 11CR entry or ATR release. To avoid severely disrupting the overall folding of the TM bundle, we chose to mutate residues (F208, F212, L266, A269, G270, F273) whose side chains do not protrude into the TM interface. We heterologously expressed these mutants in HEK cells, immunopurified the receptors, reconstituted them into lipidic bicelles, and measured the 11CR entry kinetics as well as the ATR release kinetics using the resonance energy transfer (RET) assays as described previously (Figure 2A) (Farrens et al., 1995; Schafer et al., 2016; Tian et al., 2017a, b). Briefly, 11CR entry kinetics was measured by monitoring the quenching of either the intrinsic tryptophan (Trp) fluorescence (Figure 2B, E), or an extrinsic fluorescent reporter Alexa488 (Figure 2B, F). ATR release kinetics was measured using a process that was essentially the inverse of Trp fluorescence quenching (Farrens and Khorana, 1995). For the Alexa488-based assays, we first substituted a residue in the second intracellular loop of Rho (S144) with an unnatural amino acid *p*-azido-L-phenylalanine (azF), then attached Alexa488 specifically to this site with a bioorthogonal chemistry (Figure 2C) (Tian et al., 2014; Tian et al., 2017a). These two RET-assays are functionally complementary: the Trp-based assay reports formation and dissociation of the opsin-retinal complex, while the Alexa488-based assay reports the formation of a mature pigment with the PSB characteristic for dark-state Rho (Figure 2D). Previously, we have shown that S144-Alexa488 Rho exhibits nearly identical retinal binding kinetics and energetics as wt Rho (Tian et al., 2017a), and that the regenerated Rho can be repeatedly photoactivated, confirming the functional integrity of this mutant (Tian et al., 2014). Although the Alexa488-based assay requires an additional modification step, Alexa488 fluorescence falls into a clean window of retinal absorbance, hence it is not limited by the inner filter effect as the Trp fluorescence (Tian et al., 2017a). The Alexa488-based assay is more suitable for measuring the retinal entry kinetics for “slow” mutants, for which much higher retinal concentrations are required to drive regeneration to completion. In this study, the Alexa488-based assay enabled us to analyze the binding kinetics of slow mutants at lower temperatures.

**Figure 1.**
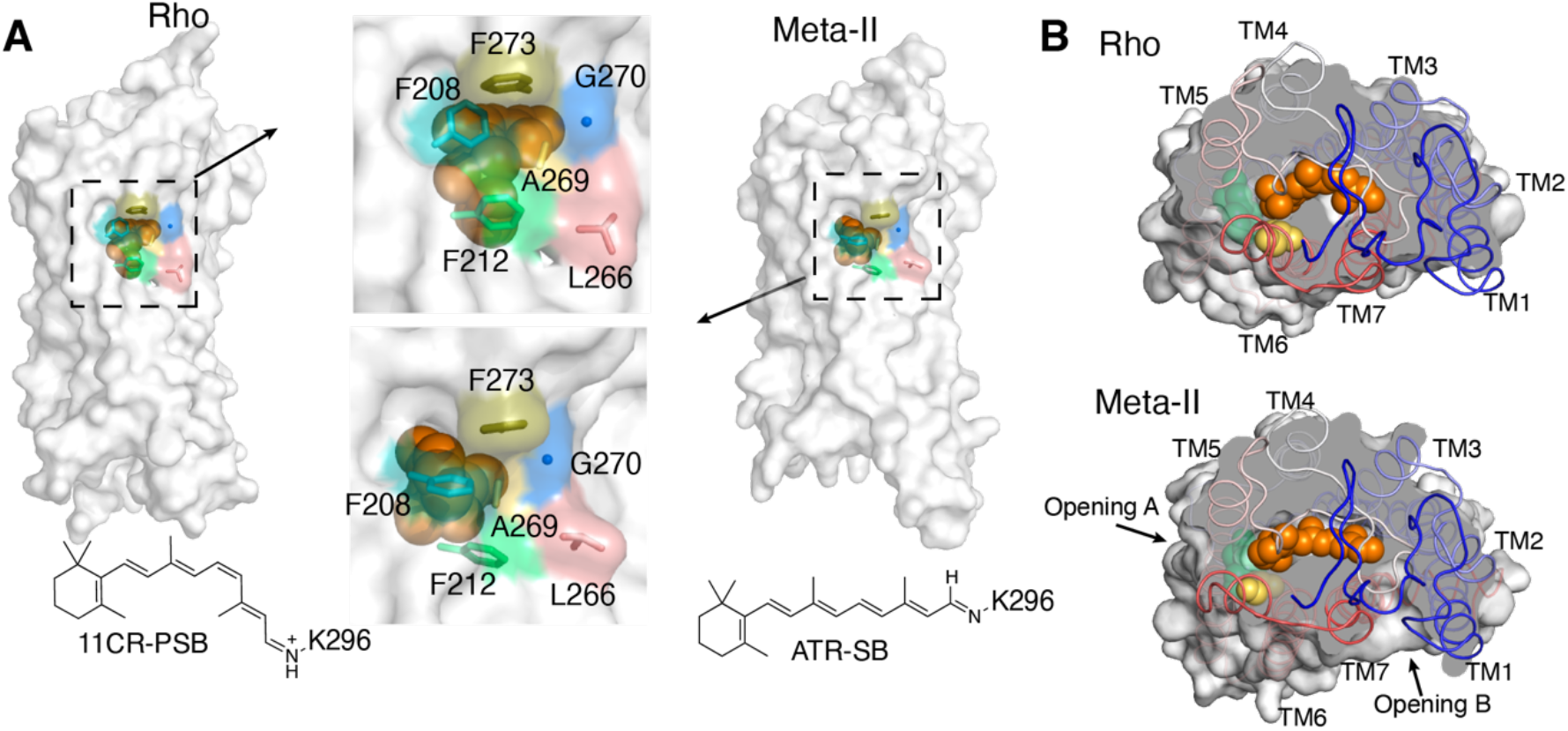
Comparing the transmembrane surfaces and ligand-binding channel of the inactive Rho and active Meta-II Rho. **A**) The side-view of 11CR-bound, inactive Rho (PDB: 1U19) and ATR-bound, active Meta-II Rho (PDB: 3PXO). Retinal is shown as spheres in *orange* and the stick and the surface representations of the mutated residues are highlighted in colors. **B**) Photoactivation causes the *cis-trans* isomerization of retinal leading to serial conformational changes, including the emergence of two openings in Meta-II Rho. The TM5/6 opening is designated as Opening A and TM1/7 opening as Opening B (retinal, *orange*; F208, *yellow*; F212, *limegreen*).

**Figure 2.**
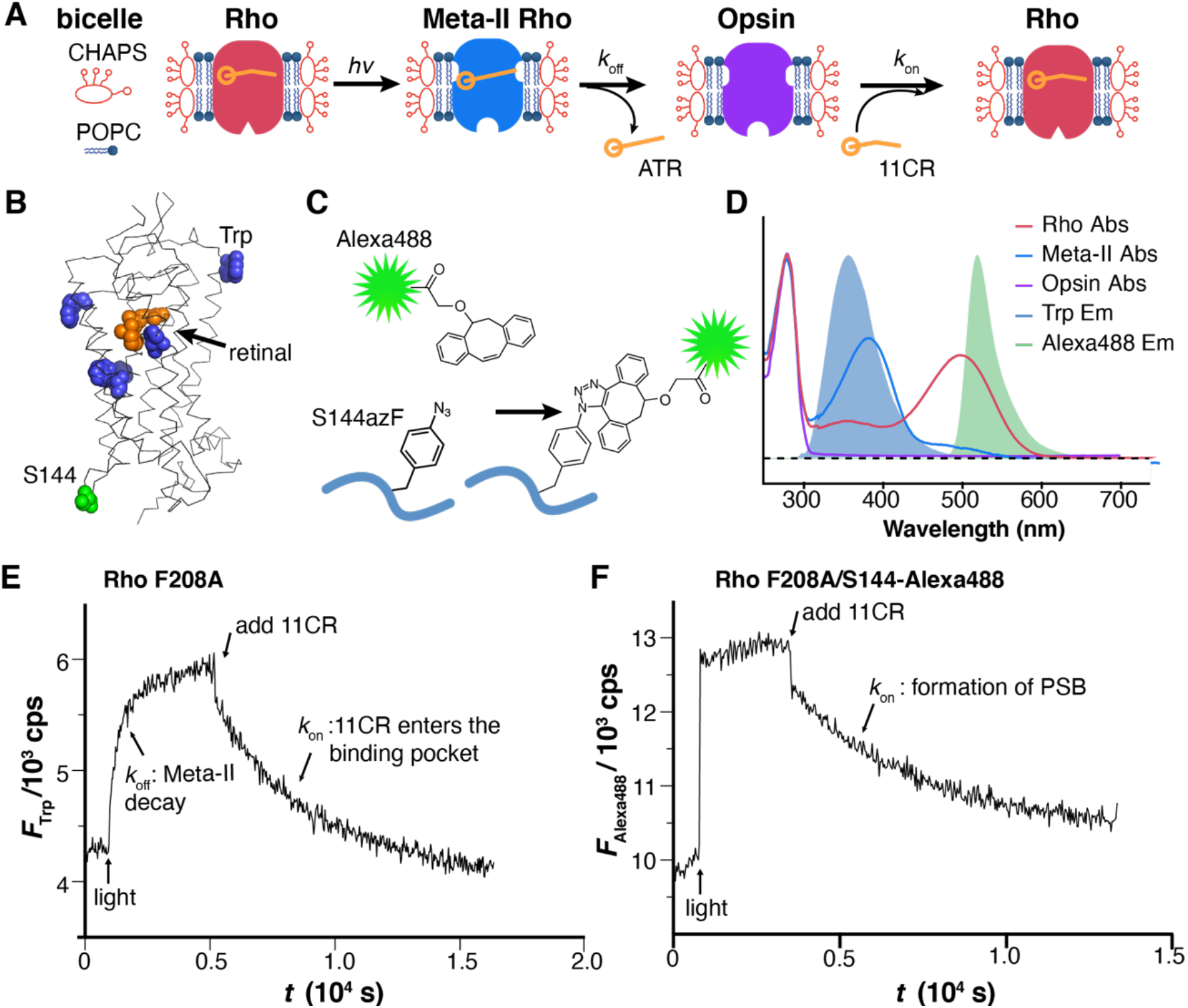
Mutational effects at putative TM5/6 11CR entry sites. **A) The assay for measuring retinal unbinding and binding kinetics.** The purified Rho is reconstituted in lipidic bicelles. **B**) The location of five Trp residues (*blue*) and S144 (*green*) relative to the 11-*cis*-retinylidene group (*orange*) in Rho (PDB: 1U19). **C**) Site-specific labeling of S144 with strain-promoted alkyne-azide cycloaddition (SpAAC). **D**) The energy transfer scheme for Trp-based and Alexa488-based assay. Note that energy transfer occurs between Trp and dark-state as well as Meta-II Rho, but is specific to dark-state Rho in the case of Alexa488. **E**) Representative trace of Trp-based RET assay. **F**) Representative trace of Alexa488-based RET assay.

We tested a panel of 24 mutants (Figure 3A, B). For Rho carrying additional mutations, the 11CR entry kinetics measurements obtained from the Trp-based and Alexa488-based assays were comparable (Figure 3A, B, Table S1). We found that mutations at the 11CR putative entry site caused a ~100-fold variation in the 11CR entry kinetics (G270L, 4.2 ± 0.4; F212W: 0.043 ± 0.022, data measured from Trp-based assay and normalized to the wt Rho), but only a modest 3-fold variation in ATR exit kinetics (F212A 2.82 ± 1.0; A269F: 0.91 ± 0.07). This 3-fold variation in ATR release kinetics resulting from TM5/6 mutations is comparable to an earlier report on zebrafish Rho (Morrow et al., 2015). The observation that the 11CR entry rate is much more affected than the ATR release rate suggested that these TM5/6 sites might be an integral part of the 11CR entry site.

**Figure 3.**
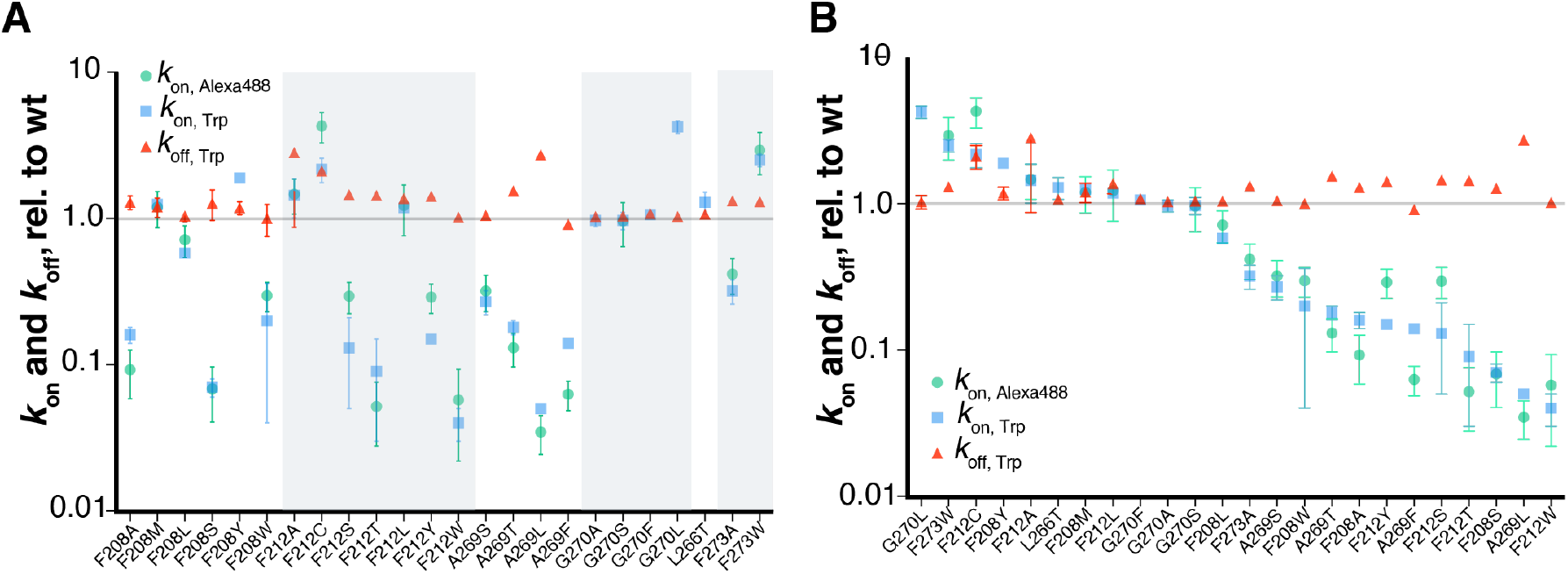
Mutational effects at TM5/6 sites. The 11CR entry kinetics is measured with Trp-based (*k*_on, Trp_) and/or Alexa488-based (*k*_on, Alexa488_) RET assays, and ATR release kinetics (*k*_off, Trp_) measured with Trp-based RET assay. **A)** Data are grouped based on the site of mutations (mean ± std). **B**) Data from **A**) shown in descending order based on *k*_on, Trp_.

Next, we asked whether the observed mutational effects were due to short-range, or potential longer-range disruptions to the ligand-binding channel. A straightforward test was to examine whether or not the mutations of residues on the opposite end of the ligand-binding channel at the pore between TM1 and TM7 behave in the mirror direction (*i.e*., whether mutating these sites cause a greater variation in the ATR release rate than in the 11CR binding rate). We tested T94I (Figure 4A) and found a moderate decrease in the 11CR entry kinetics (0.80 ± 0.04), but a significant decrease on the ATR release kinetics (0.17 ± 0.02) (Table S1). However, a caveat here is that these TM1/7 residues, being relatively juxtaposed to the SB bond with retinal, may simultaneously affect the Schiff base chemistry and the folding of the putative ligand-binding channel (Janz et al., 2004). Alternatively, the hydrophobic substitution T94I could lead to a reduction of the local concentration of water molecules available for the SB hydrolysis reaction, and it would be difficult to dissect their respective contributions to the overall mutational effect.

**Figure 4.**
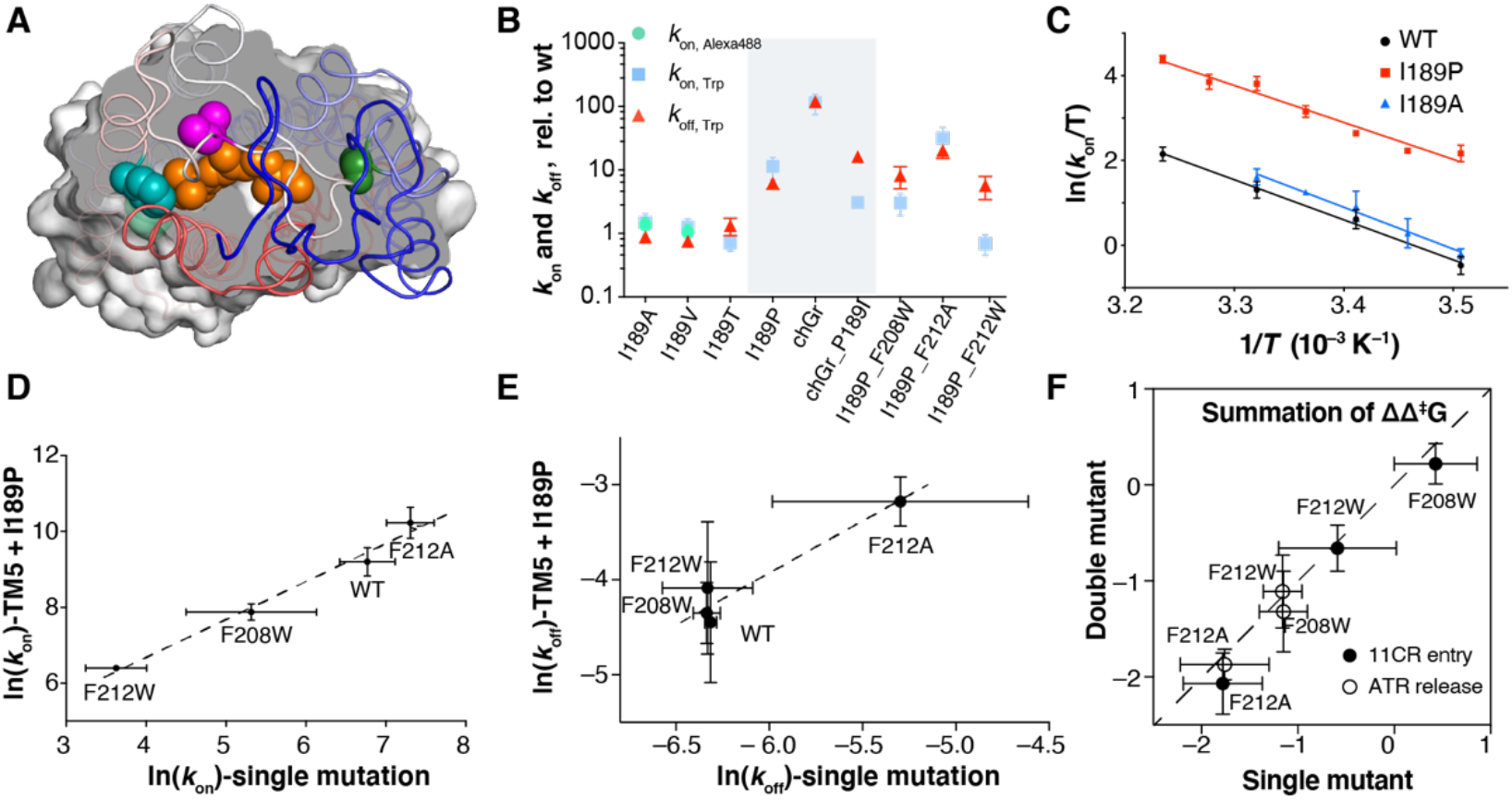
Double mutant analysis. **A)** Two TM5 sites (F208, *teal*; F212, *limegreen*), I189 (*magenta*), and T94I (*forest green*) highlighted in Rho (PDB: 1U19). **B**) Mutational effect at site 189. I189P simultaneously increases the 11CR entry and ATR release kinetics (mean ± std). **C)** The Eyring plot for I189P and I189A mutants. The kinetics (mean ± std) were measured using the Alexa488-based assay. **D**) Double logarithmic plot of *k*_on, Trp_ of TM5 single mutant versus TM5/I189P double mutants. I189P reduces the activation free energy (Δ^‡^*G*) by 1.6 ± 0.3 kcal mol^-1^. The fitting equation is ln(*k*_on_(I189P)) = (0.997 ± 0.093)×ln(*k*_on_(I189)) + (2.69 ± 0.55) (*R*^2^ = 0.98). **E**) Logarithmic plot of *k*_off_ Trp in TM5 single mutant and TM5/I189P double mutants. I189P reduces the activation free energy (Δ^‡^*G*) by 1.6 ± 0.8 kcal mol^-1^. The fitting equation is ln(*k*_on_(I189P)) = (1.09 ± 0.23)×ln(*k*_on_(I189)) + (2.6± 1.4) (*R*^2^ = 0.92). **F**) Summation of ΔΔ^‡^*G* in single mutants vs. ΔΔ^‡^*G* in the corresponding double mutants.

We reasoned that a better approach would be to test a site that alters the middle segment of the ligand-binding channel. We therefore chose to examine I189, a conserved molecular determinant that distinguishes between rod and cone opsins. In rod receptors, site 189 is invariably an Ile or Val, while in cone receptors a Pro is very highly conserved (Ebrey et al., 2001). One exception is lamprey rhodopsin, which is an evolutionary intermediate between cones and rods and carries P189 (Yanagawa et al., 2015). In Rho, I189 is located at the end of a β-hairpin formed by the second extracellular loop that functions as the “cap” of the ligand-binding channel (Figure 4A). The I189P mutation in Rho has been reported to increase the Meta-II decay rate (Kuwayama et al., 2002), increase the regeneration rate (Imai et al., 2005), as well as elevate the thermal isomerization rate of 11CR to ATR (Yanagawa et al., 2015).

Among the I189 mutants tested here (Figure 4B), we found that the reduced size of the aliphatic side chain (I189A, I189V) only slightly accelerated 11CR entry kinetics and reduced the ATR release rate. Eyring plots of the kinetics data showed that I189A only made a slight difference to the transition energy (Figure 4C). We also observed a decreased thermal stability for I189A at higher temperature (unpublished data). Substitution of an aliphatic Ile with a polar Thr moderately slowed down 11CR entry and accelerated ATR release. Overall, the effects of the side chain size and polarity are modest. In contrast, we observed that the I189P mutation dramatically increased both 11CR entry and ATR release rate (Figure 4B), which is consistent with previous reports (Imai et al., 2005; Kuwayama et al., 2002). Back-converting the Pro in a chicken green cone opsin to Ile significantly slowed down 11CR binding and ATR release. Eyring analysis indicated that the I189P substitution accelerated 11CR entry by increasing the entropic factor. These data suggest that I189P significantly eases the structural constraints on retinal movement within the channel, possibly through proline’s ability to introduce kinks into the peptide backbone.

It was well established that the summation of the energetic effects of single mutations is nearly equal to the free energy change measured in the corresponding multiple mutants (Dill, 1997; Wells, 1990). Deviation from this simple additivity rule is a strong indicator that 1) the mutated residues are interacting with each other directly or indirectly, or 2) the mutation causes a change in mechanism or rate-limiting step. Therefore, we measured the 11CR entry and ATR release rates for three double mutants consisting of a TM5 mutation and I189P. We found that the I189P mutation and TM5/6 mutations on 11CR entry rate were highly energetically additive (Figure 4D-F). In the natural logarithmic plot of the 11CR entry and ATR release kinetics, the slope was fitted to 1.00 ± 0.09 (R^2^ = 0.98) and 1.1 ± 0.2 (R^2^ = 0.92), respectively (Figure 4D, E).

We calculated the change of activation free energy (ΔΔ^‡^*G*) for these mutants. We found that the summation of ΔΔ^‡^*G* for single mutants is close to the ΔΔ^‡^*G* of corresponding double mutants (Figure 4F). I189P reduced Δ^‡^*G*_on_ by 1.6 ± 0.3 kcal mol^-1^ and Δ^‡^*G*_off_ by 1.6 ± 0.8 kcal mol^-1^. This additivity shows that I189 and the TM5/6 site are non-interacting structural determinants for the retinal-binding channel. Intriguingly, the changes of Δ^‡^*G*_on_ and Δ^‡^*G*_off_ are statistically the same, suggesting that I189P reduces constriction for the retinal ligands to traverse through the ligand-binding pathway in either direction. The lack of synergistic effects between I189P and TM5 mutations suggest that the observed mutational effects at the TM5/6 channel is localized within a short-range. Thus, the most parsimonious explanation for the observed large variation in 11CR entry kinetics is that the TM5/6 channel constitutes the 11CR entry site.

We further analyzed mutations at three TM5/6 sites based on their side chain properties (Figure 5). We tested polar mutations at three sites (F208, F212, A269). Ser and Tyr differ only from Ala and Phe in having additional hydroxyl groups, and the comparison of Ser/Ala and Tyr/Phe pairs are particularly meaningful as these substitutions involve minimal steric changes apart from the hydroxyl group. We also included Thr, an analogue of Ser. We found all the polar mutations, except for F208Y, reduced the 11CR entry kinetics (Figure 5A). We could not measure the retinal entry kinetics for A269Y because this mutant simply could not be regenerated even after prolonged incubation with a high concentration of 11CR (15 μM, over 48 h) (*i.e*., 11CR entry for A269Y is either indefinitely slow or the mutation irreversibly denatures the receptor).

**Figure 5.**
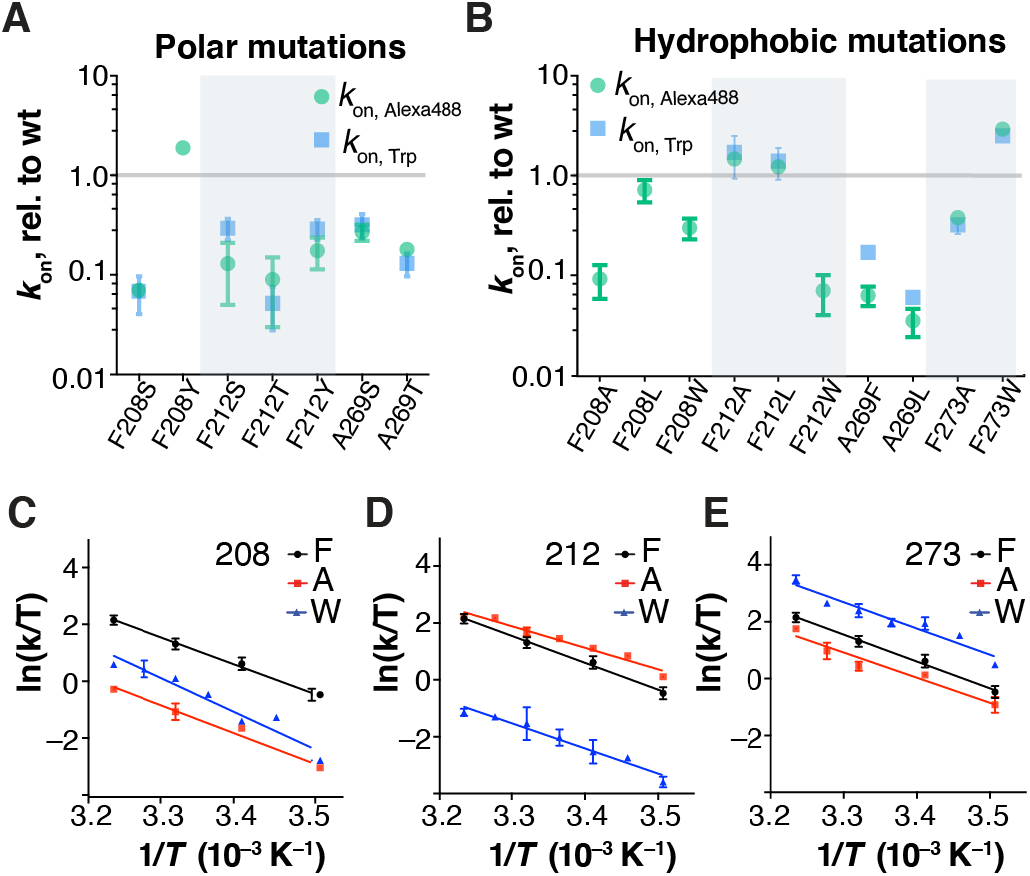
Comparison of the effects of polar and hydrophobic mutations on 11CR entry kinetics. **A)** The effect of polar mutations on 11CR entry kinetics (mean ± std). **B)** The effect of hydrophobic substitutions on 11CR entry kinetics (mean ± std). The A269W mutant only expressed at very low level and failed to regenerate. **C-E**) The Eyring plots of the 11CR entry kinetics for the following mutants: F208A/W, F212A/W, F273A/W. The kinetics were measured using the Alexa488-based assay.

The data for hydrophobic mutations revealed a more complex pattern (Figure 5B). We varied the steric hindrance at these sites by installing a series of hydrophobic substitutions (Ala, Phe, Leu, Trp). For F212 and A269, 11CR entry kinetics is negatively correlated with the size of the side chain. The F273 mutations are the opposite of that of F212 and A269; decreasing the side-chain size led to slower retinal binding. For mutations at F208, increasing the side-chain size (F208W) or decreasing it (F208A) both resulted in slower 11CR entry kinetics.

We further measured the temperature dependence of 11CR entry kinetics for six mutants, F208A/W, F212A/W, and F273A/W using the Alexa488-based 11CR entry assay (Figure 5D-F). All of the Eyring plots for the wt and the mutants showed a high degree of linearity (R^2^ = 0.92~0.97). Two mutations, F208W and F212A, caused significant change in the activation enthalpy. F208W increased the activation enthalpy (Δ^‡^*H*) by 4.8 ± 3.1 kcal mol^-1^, while F212A reduced it by 3.7±1.6 kcal mol^-1^. The thermodynamic effects of F273A/W are less pronounced compared with F208A/W and F212A/W.

## DISCUSSION

### Experimental design

An earlier study on retinal entry kinetics reported non-localized mutational effects and did not pinpoint a putative entry site (Piechnick et al., 2012). We contend that the design of the preceding study was limited in two important ways. First, if the side chain of a residue protrudes into the TM domain, mutating this site can disrupt the overall receptor folding. As a result, the mutational effect is more likely to be non-localized. In the present study, we chose to focus on residues whose side chains faced outward from the TM region. Second, for ATR release, it is well established that hydroxylamine, which cleaves the SB bond, accelerates Meta-II decay in both wt Rho and mutants by about two-orders of magnitude (Piechnick et al., 2012), suggesting that SB hydrolysis could be a rate-limiting step of ATR release. The implication for this mutagenesis study is that a mutation affecting SB chemistry is likely to cause a greater change in the apparent ATR release rate than a mutation affecting the channel property. Since residues at TM1/7 are prone to affect both SB chemistry and folding of the channel, the mutational effects at these sites are difficult to interpret. Thus, we focused on the TM5/6 end of the ligand-binding channel and only mutated residues whose side chains face outward.

RET-based assays have been widely used to study intra- and inter-molecular dynamics. When it comes to studying ligand binding, a challenge is to find the appropriate energy transfer partner with the ligand. In the case of Rho, the intrinsic Trp fluorescence serves as a good reporter for retinal entry, except for the slower mutants. However, in most cases, Trp fluorescence has insufficient spectral overlapping with the ligand. Our bioorthogonal labeling strategy represents an attractive alternate: since the labeling is modular, site-specific, and the fluorophore can be chosen with flexibility, RET-assays can be developed to probing ligand binding in a variety of GPCR-ligand pairs. Moreover, genetic code expansion eliminates the need to generate and validate single-accessible Cys mutant that is often required by the classic site-specific protein labeling methods.

### The permissive opsin conformation for 11CR binding

In the dark-state Rho structure, the inactive opsin-11CR covalent complex (Li et al., 2004; Okada et al., 2004; Palczewski et al., 2000) features an enclosed ligand channel. The electron density between TM5 and TM6 is slightly delocalized at the position of F212 and A269 as a result of the kink in TM6. Upon photoactivation, the cytoplasmic end of TM6 moves outward, creating obvious openings to the lipidic membrane at each end of the ligand channel, one between TM1 and 7, the other between TM5 and 6 (Choe et al., 2011; Standfuss et al., 2011). Previously it was proposed that 11CR binding with opsin requires the active receptor conformation, and that retinal enters and exits through either or both of these two openings (Hildebrand et al., 2009; Piechnick et al., 2012).

However, several lines of new experimental observations contradict this “active conformation” model. First, we have shown previously that the opsin reconstituted in the lipidic bilayer of bicelles robustly recombines with 11CR one hour after photoactivation, while opsin in DDM failed to bind with 11CR in a temperature-dependent manner (Tian et al., 2017a). An EPR (electron paramagnetic resonance) DEER spectroscopy study indicated that the cytoplasmic end of TM6 reset spontaneously within 30 min in nanodiscs, but it failed to do so in DDM micelles (Van Eps et al., 2017). In fact, the 11CR entry assay performed for DDM-solubilized Rho typically carries a N2C/D282C stabilizing mutation (Piechnick et al., 2012; Schafer and Farrens, 2015). It would be interesting to test whether this additional disulfide bond promotes TM6 resetting in detergents. Second, a constitutively activating mutation M257Y and transducin-mimic peptide fused to the C-terminal tail of Rho, both shifting the equilibrium of the receptor population towards the active state (Deupi et al., 2012; Han et al., 1998; Tsukamoto et al., 2013), selects for ATR rather than 11CR (Schafer and Farrens, 2015). These independent lines of experimental evidence together predict that the permissive opsin conformer for 11CR entry (Ops**) is characterized by TM6 resetting, rather than outward movement of TM6, *i.e*., it shows greater resemblance to the inactive conformation than to the active conformation.

### An allosteric model for 11CR entry

Our data show that 1) mutating the TM5/6 end of the ligand-binding channel causes a much greater change to the 11CR entry kinetics than to the ATR release kinetics, and 2) these mutational effects on the TM5/6 pore are localized. These observations strongly support a scenario that an entry site between TM5/6 exists to allow 11CR to access the ligand-binding channel, and that this entry site roughly co-localizes with the TM5/6 opening observed in the Meta-II Rho. How does this scenario reconcile with our earlier statement that Ops** is likely to resemble the inactive conformation, given the knowledge that TM region of the inactive conformer is completely enclosed? Is it possible for Ops** to be simultaneously characterized with TM6 resetting and an accessible opening?

Another question arising from our data is how to rationalize the mutational effects of hydrophobic substitutions? We initially made an intuitive prediction – mutational effects at the putative entry site would be dominated by the steric factor, (*i.e*., larger side chains should slow down 11CR entry, while smaller side chains should accelerate it). We indeed found two sites, F212 and A269, that conformed to this predication. The consistent trend observed for the polar mutations at F212 and A269 further suggests that these two sites act in concert. On the other hand, we also observed residues F208, F273, and G270L deviate from the predicted pattern. F208A, F208W, and F273A slowed down 11CR entry while F273A and G270L accelerated it. In fact, the Farrens group has reported similar observations for F208A and F273A (Schafer and Farrens, 2015). They further speculated the making the larger pore would result in less efficient “trapping” of 11CR inside the channel, thus explaining the slower 11CR entry kinetics. However, our data on F208W and F212W seems to refute this hypothesis, unless the “trap” has very specific steric requirement and the larger Trp residue prevents the “trap” to close properly.

Previous experiments that sought to “modify the size of opening” are implicitly based on the concept of “conformational selection”: a conformational change occurs *prior to* the binding event, and ligand stabilize the permissive, typically high-energy, conformation for binding. The role of conformational selection is widely appreciated for ligand in Rho (Schafer and Farrens, 2015; Schafer et al., 2016) as well as a wide range of GPCRs (Latorraca et al., 2017). The “conformational selection” model takes a static view of the ligand binding channel. In the case of Rho regeneration, the underlying assumptions were: 1) the size of the 11CR entry opening is sufficiently large to accommodate the cross section of 11CR, including the β-ionone ring; 2) the size of the opening remains roughly constant over the course of ligand entry; 3) the entry opening must promptly close right after ligand entry to protect the PSB. In this picture, installing a larger side chain should invariably decrease the 11CR entry kinetics, and *vice versa*.

Here, we propose an “allosteric opening” model for 11CR entry (Figure 6). The pore opening does not have to be sufficient large in the very beginning. Rather, this opening is dynamically reformed as 11CR traverses through the channel, possibly accompanied by additional conformational change. Without the inverse agonist 11CR stabilizing the inactive state, the apoprotein opsin alone is likely to be more flexible and dynamic than Rho. Single-molecule fluorescence experiments (Maeda et al., 2018) and electrophysiological recordings (Sato et al., 2019) have confirmed the existence of multiple conformers for opsin (Oprian et al., 1987). This flexibility of the opsin backbone makes it conceivable for 11CR to force open TM5/6. In fact, the notion of dynamically-adjusted ligand-binding channels has been suggested by molecular dynamic simulations on β_2_ adrenergic receptors (Dror et al., 2011) and human protease-activated receptor 1(Zhang et al., 2012).

**Figure 6.**
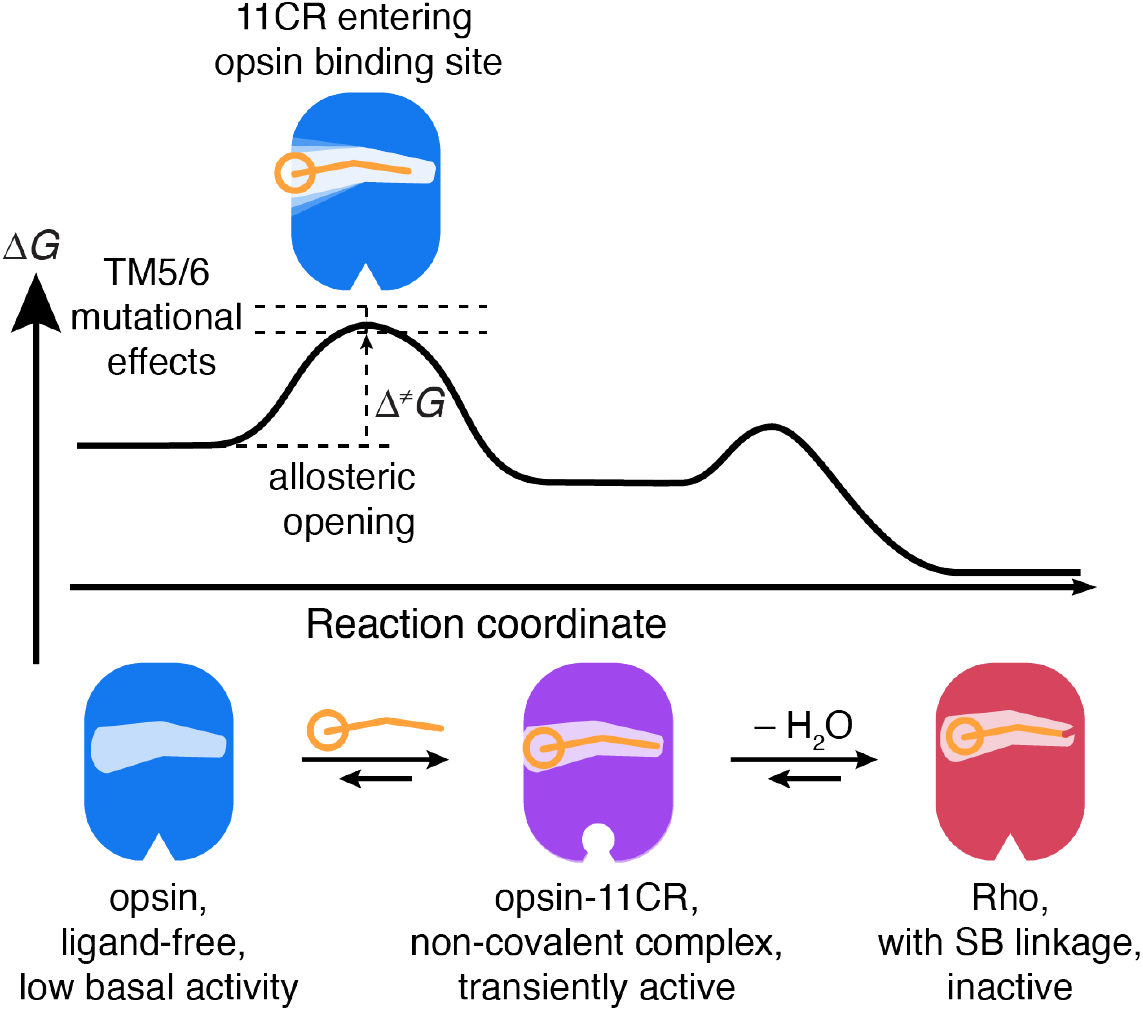
A model for 11CR entry in opsin.

The energetic cost for reforming the opening might underlie the notably high activation barrier (Δ^‡^*G*_on_ = 13.5 kcal mol^-1^) for 11CR binding with opsin (Tian et al., 2017b). By comparison, Δ^‡^*G*_on_ for high affinity ligands for β_2_ adrenergic receptor (Rasmussen et al., 2011; Rosenbaum et al., 2007) is lower by 3 kcal mol^-1^. The extraordinarily high energy barrier may arise from the additional conformational change of the protein backbone required for 11CR entry, as well as the energetic cost for Schiff base formation.

The general trend for polar mutations supports our hypothesis that TM5/6 is the 11CR entry site. In such a scenario, the aldehyde group should precede the polyene chain and the β-ionone ring into the channel. A plausible mechanism is that the carbonyl group of retinal may interact with the substituted hydroxyl group through dipole-dipole interaction or hydrogen bonding to form a transiently stabilized intermediate, resulting in slower binding kinetics. Alternately, these additional polar groups in the low dielectric membrane environment may be forced to hydrogen-bond with nearby polar residues and thus limit the flexibility of the entry site to undergo the necessary conformational change.

Importantly, our model may rationalize the counterintuitive mutational effects of F208A/W, F273A/W and G270L. While steric hindrance is necessarily localized, localized effect is not limited to steric hindrance. A mutation not only changes the packing density *in situ*, but also affects any interaction with nearby residues and the incoming 11CR. As a result, steric hindrance alone may not dominate the network effect of a mutation on 11CR binding kinetics; the altered energetic cost of opening expansion must be taken into account. Additional molecular dynamics stimulation may be needed to explore how these mutations alter the energetic landscape of ligand channel.

Our model is also consistent with the electrophysiological observation that 11CR binding causes transient activation of opsin (Kefalov et al., 2001). The change of receptor signaling state may be explained by major conformational change necessitated by 11CR entry. Here, the transiently active non-covalent complex of 11CR and opsin arises when retinal is sitting fully in the binding pocket, but due to the lack of a Schiff-base between 11CR and K296, the additional oxygen atom (from the retinal) increases the distance between helix 7 and the beta ionone ring. That increase in distance is not unlike the stretching from photoisomerization of 11-*cis* to all-*trans* retinylidene causing formation of the active state. Therefore, the longer distance in the non-covalent complex activates the receptor and once the Schiff base forms, the receptor resets to the inactive state. The non-covalent state is an intermediate state after the barrier has crossed and before the inactive state has been reached (Figure 6). As we do not observe any measurable delays between retinal entry as measured in the Trp quenching experiment and the formation of the PSB as measured by the Alexa-488 quenching experiment, the barrier between this intermediate and the inactive state should be smaller than the barrier determined by the transition state involving the widening of the TM5/6 opening, which is the rate limiting step of the regeneration reaction. We have previously shown that the overall energy diagram of the ligand binding reaction in rhodopsin is comparable to a GPCR with a tight-binding, non-covalent ligand (Tian et al., 2017b). The Schiff-base contributes only a small amount of the barrier that has to be crossed upon ligand release.

It is worth noting that our “allosteric” model and the “conformational selection” model are not mutually exclusive. To dissect their respective contributions, it is important to elucidate the initial state of the opening for 11CR immediately before the ligand binding process. An entry pore that requires weakening of electron density between TM5 and 6 may arise from one of, or a combination of the following plausible mechanisms: 1) the opening entirely emerge as a result of the interaction between the incoming 11CR and opsin; 2) a small population of Ops** preexists in equilibrium as suggested by Farrens et al. (Schafer and Farrens, 2015); 3) the real structure of inactive opsin is in fact different from the apoprotein in the Meta-II structure and the inactive opsin itself contains such an entry site and qualifies as Ops**. Our data cannot formally distinguish among these possibilities. In future studies, NMR should be utilized to elucidate the conformational dynamics of opsin in the course of 11CR entry. In particular, the sites of interest identified in this study could be replaced with NMR-active unnatural amino acids to probe the conformational dynamics (Jones et al., 2010).

## CONCLUSION

Accumulating structural studies of GPCRs reveal a lateral binding pocket that is inaccessible from the extracellular milieu (Szlenk et al., 2019). While intermembranous ligand binding pathways for hydrophobic ligands appears plausible, current understanding of binding pathways in GPCRs is primarily derived from computational simulations (Alfonso-Prieto et al., 2019; Ibrahim et al., 2019; Marino et al., 2018; Stanley et al., 2016). Direct experimental evidence on intermembranous ligand entry sites is rare (Bokoch et al., 2018; Piechnick et al., 2012). Here we addressed the question of native ligand (11CR) entry in the visual photoreceptor Rho, a prototypical GPCR. Through the engineering of a sensitive RET sensor, improved mutagenesis design, and extensive energetic analysis, we were able to show that amino acid residues close to a putative pore between TM5 and TM6 display the expected characteristics of a ligand entry site. Mutating these sites caused a much greater variation in the 11CR entry kinetics compared with the ATR release kinetics. The effects of the side chain substitutions lining the pore were localized. Based on our own data and the existing literature, we propose that 11CR entry involves an allosteric opening of a pore in a permissive opsin conformation, which dynamically adjusts to the incoming 11CR. Our findings highlight the importance of understanding how conformational dynamics regulates ligand binding in a GPCR. Moreover, our approach of engineering site-specific RET reporter and double-mutant energetic analysis can be extended to study ligand binding in a wide range of intermolecular interactions.

## EXPERIMENTAL SECTION

### Materials

*E. Coli* TOP10 cells (Invitrogen) were used for plasmid propagation and isolation. 1D4-Sepharose 2B was prepared from 1D4 mAb and CNBr-activated Sepharose 2B (2 mg IgG per mL packed beads) as described (Knepp et al., 2011; Oprian et al., 1987). Alexa488-DIBO was obtained from Molecular Probes/Thermo Fisher Scientific as a dry powder, dissolved in DMSO (5 mM), and stored at −20 °C. n-dodecyl-β-d-maltoside (DDM) was obtained from Anatrace. 1-palmitoyl-2-oleoyl-sn-glycero-3-phosphocholine (POPC) was obtained from Avanti. Plasmids for amber codon suppression were reported in a previous publication (Ye et al., 2008). Mutations were introduced into bovine rhodopsin using the QuikChange site-directed mutagenesis kit (Agilent) according to the manufacturer’s protocol.

### Heterologous expression of wt and mutant Rho in mammalian cell culture

The wt and azF-tagged Rho were expressed in HEK293F suspension cell culture. The HEK293F suspension cells were cultured in serum-free FreeStyle 293 expression medium in a 125-mL disposable, sterile, polycarbonate Erlenmeyer flask (Corning) at 37°C in 5% CO_2_ atmosphere and constantly shaked at 125 rpm. Transfection of suspension cells was done using FreeStyleMax according to the manufacturer’s protocol. For a 30-mL culture, 38.6 μg of plasmid DNA (for amber codon suppression 18.4 μg of pMT4.Rho containing the amber codon, 18.4 μg of pSVB.Yam, and 1.84 μg pcDNA.RS-azF were mixed together) was used. For wt and Rho mutants without unnatural amino acids, cells were harvested 48 hours post-transfection. For azF-tagged Rho mutants, FreeStyle 293 expression medium was supplemented with 1 mM azF and the cells were harvested 96 hours post-transfection. The total cell number upon harvesting normally ranged from 6×10^7^ to 8×10^7^. The harvested cells were resuspended in DPBS (Gibco, supplemented with leupeptin and aprotinin, Sigma) at a density of 10^7^ cells/mL in a 15-mL conical, polypropylene tube (Falcon). In the dark room, 11-*cis*-retinal (1.4 mM ethanol solution) was added into the cell suspension to a final concentration of 5 μM. (Oprian et al., 1987; Starace et al., 1998). The suspension was nutated at 4°C at least overnight. Excess 11-*cis*-retinal was removed by spinning down the cells and discarding the supernatant fraction. The regenerated cells can be immediately used, or may be stored at −20°C for several months.

### Immunopurification

The 11-*cis*-retinal regenerated cells expressing Rho mutants were lysed with the solubilization buffer (1 mL per 10^7^ cells, 1% (w/v) DDM, 50 mM HEPES or Tris-HCl, pH 6.8, 100 mM NaCl, 1 mM CaCl2 with Complete EDTA-free Protease Inhibitor Cocktail, Roche) for at least 1 h at 4°C. The lysate was cleared by centrifugation at 100,000×g for 30 min and incubated overnight at 4°C with 1D4-mAb-sepharose 2B (100 μL). The resin was transferred into a 1.5-mL Eppendorf tube and washed three times for 30 min each with 0.5 mL reaction buffer (0.1% DDM in DPBS, pH 7.2). AzF-tagged mutants was labeled before the elution step. Non-azF-tagged mutants was directly eluted with a competitive peptide.

### Bioorthogonal labeling

Then the reaction buffer (200 μL) was mixed with the resin (100 μL) to give 300-μL slurry. The Alexa488-DIBO stock solution (5 mM in DMSO) was directly added to into the reaction mixture to give the appropriate final concentration. The reaction was agitated with a thermomixer at 25 °C. For labeling S144azF, we used 50 μM of Alexa488-DIBO and allowed the reaction to proceed for 18 hours. The reaction was stopped by centrifugation and removal of the supernatant fraction. The resin was then transferred into a microporous centrifugal filtering unit (Microcon-MC pore size 0.48 μm, Millipore). The resin was first washed with the reaction/wash buffer for three times (30 min incubation each time) to deplete the unreacted dyes, and then with a low-salt buffer (0.1% (w/v) DDM, 2 mM sodium phosphate buffer, pH 6.0). The receptor was eluted with elution buffer (100 μL, no less than the volume of the resin; 0.33 mg/mL C9 peptide in 0.1% (w/v) DDM, 2 mM sodium phosphate buffer, pH 6.0). The resin was incubated with the elution buffer on ice for at least 1 h. The purified receptor was collected in a clean 1.5-mL Eppendorf tube. The elution was repeated a second time. The combined elutions were supplemented with 150 mM NaCl and characterized by UV–Vis spectroscopy. Purified samples were stored at −80°C. The yield from 10^7^ HEK293F cells was typically 0.5–1 μg Alexa488-labeled azF-Rho (Tian et al., 2014).

### Reconstitution of purified Rho mutants into POPC/CHAPS bicelles

The POPC lipids were dissolved in 20% (w/v) CHAPS solution at 1:1 POPC-to-CHAPS ratio (w/w) using repeated freeze-thaw cycles and diluted with water to make a 10% (w/v) stock bicelle solution. The 10% stock solution was aliquoted, kept at −20°C and thawed prior to use. The final working solution (1% (w/v) POPC, 1% (w/v) CHAPS, 125 mM KCl, 25 mM MES, 25 mM HEPES, 12.5 mM KOH, pH 6.0) was prepared freshly by diluting 10% POPC/CHAPS stock solution with the corresponding concentrated buffer stocks. At the beginning of the retinal binding assay, the frozen purified Rho samples (30 μL) were thawed on ice and diluted into the freshly prepared bicelle buffer (450 μL).

### Measurement of 11CR entry kinetics or ATR release kinetics based on quenching of Trp or Alexa488 fluorescence

All of the 11CR and ATR release assays were performed in 1% POPC/CHAPS bicelle solutions (10 mg/mL POPC and 10 mg/mL CHAPS). Fluorescence measurement was done on a SPEX Fluorolog tau-311 spectrofluorometer (Horiba) in photon-counting mode. The temperature was controlled by circulating pre-heated or pre-chilled water through the recording chamber. 30 μL of purified Rho was added into 450-μL of POPC/CHAPS buffer. In the Trp-based fluorescence experiment, the excitation wavelength was 295 nm with a 0.6-nm band-pass, and the emission was measured at 330 nm with a 15-nm band-pass. The concentration of Rho was typically 0.25-0.30 μM. In the Alexa488-based experiment, the sample was excited at 488 nm with a 0.2-nm band-pass, and the emission was measured at 520 nm with a 15-nm band-pass. The concentrations of Alexa488-Rho in these experiments were lower, normally in the range of 5-50 nM. The sample was constantly stirred to facilitate mixing. The fluorescence signal was integrated for 2 s in 30-s intervals. For each measurement, 11CR ethanolic stock solution was freshly diluted in POPC/CHAPS bicelle buffer and used within 12 hours. The actual concentration of 11CR was determined by UV-vis spectroscopy using the extinction coefficient of 11CR in POPC/CHAPS bicelles (ε378_nm_=25,600 M^-1^ s^-1^) (Tian et al., 2017a). 20 μL of 11CR working dilution was added to the cuvette to give a final concentration in the range of 0.5 to 30 μM, adjusted based on the first measurement of 11CR entry kinetics. The decrease of Trp or Alexa488 fluorescence was fitted with a pseudo first-order exponential decay model to derive the apparent regeneration rate (*k*_obs_). The second-order rate constant (*k*_2_) for the recombination reaction between opsin and retinal was calculated as *k*_2_ = *k*obs /[retinal]. The full details of preparing POPC/CHAPS buffer and retinal concentration measurement have been reported in an earlier publication (Tian et al., 2017a).

### Data Availability Statement

The 11CR binding and ATR unbinding kinetics are provided in the Supplemental Information.

## ACKNOWLEDGMENTS

We acknowledge the generous support from the Crowley Family Fund and the Danica Foundation. This work was also been generously supported by an International Research Alliance with Professor Thue W. Schwartz at The Novo Nordisk Foundation Center for Basic Metabolic Research (http://www.metabol.ku.dk) through an unconditional grant from the Novo Nordisk Foundation to University of Copenhagen. We also acknowledge the support from NIH R01 EY012049 (T.P.S. and T.H.) as well as the Tri-Institutional Training Program in Chemical Biology in supporting H.T.

## AUTHOR CONTRIBUTIONS

H.T., T.P.S., and T.H designed the study, analyzed the results, and wrote the manuscript. H.T., T.H., K.M.G. and M.K. conducted the experiments.

The authors declare no conflict of interest.

## SUPPLEMENTAL INFORMATION AVAILABLE

### ABBREVIATIONS

11CR: 11-*cis*-retinal
ATR: all-*trans*-retinal
azF: *p*-azido-L-phenylalanine
CHAPS: 3-[(3-cholamidopropyl)-dimethylammonio]-1-propanesulfonate
DIBO: dibenzocyclooctyne
DDM: *n*-dodecyl-β-D-maltoside
GPCR: G protein-coupled receptor
POPC: 1-palmitoyl-2-oleoyl-*sn*-glycero-3-phosphocholine
PSB: protonated Schiff base
RET: resonance energy transfer
Rho: rhodopsin
ROS: rod outer segment
SB: Schiff base
SpAAC: strain-promoted [3+2] azide-alkyne cycloaddition
Trp: tryptophan
UAA: unnatural amino acid
wt: wild-type.

## Notes

### Competing Interest Statement

The authors have declared no competing interest.

