## Supplemental Information for "Entry Pathway for the Inverse Agonist Ligand in the G Protein-Coupled Receptor Rhodopsin"

**Table S1. The kinetics of 11CR binding and ATR unbinding measured at 28 °C**

| | $k_{on,\ Alexa488}$ ($\times 10^3\ M^{-1}\ s^{-1}$) | $k_{on,\ Trp}$ ($\times 10^3\ M^{-1}\ s^{-1}$) | $k_{off,\ Trp}$ ($\times\ 0.001\ s^{-1}$) |
|---|---|---|---|
| wt | 1.12 ± 0.25 | 1.03 ± 0.04 | 1.77 ± 0.13 |
| F208A | 0.103± 0.030 | 0.170 ± 0.019 | 2.28 ± 0.18 |
| F208M | 1.34 ± 0.22 | 1.30 ± 0.04 | 2.12 ± 0.27 |
| F208L | 0.802 ± 0.086 | 0.596 ± 0.016 | 1.84 ± 0.07 |
| F208S | 0.077 ± 0.026 | 0.073 ± 0.012 | 2.25 ± 0.51 |
| F208Y | N.A. | 1.95 ± 0.05 | 2.08 ± 0.15 |
| F208W | 0.333 ± 0.016 | 0.20 ± 0.17 | 1.78 ± 0.43 |
| F212A | 1.64 ± 0.25 | 1.49 ± 0.44 | 5.0 ± 3.4 |
| F212C | 4.80 ± 0.33 | 2.25 ± 0.41 | 3.74 ± 0.63 |
| F212S | 0.330 ± 0.032 | 0.136 ± 0.077 | 2.57 ± 0.17 |
| F212T | 0.058 ± 0.024 | 0.098 ± 0.058 | 2.56 ± 0.98 |
| F212L | 1.37 ± 0.43 | 1.22 ± 0.03 | 2.42 ± 0.30 |
| F212Y | 0.325 ± 0.016 | 0.152 ± 0.006 | 2.52 ± 0.03 |
| F212W | 0.064 ± 0.037 | 0.38 ± 0.014 | 1.81 ± 0.06 |
| A269S | 0.358 ± 0.059 | 0.281 ± 0.048 | 1.86 ± 0.09 |
| A269T | 0.146 ± 0.19 | 0.191 ± 0.020 | 2.73 |

| | | | |
|---|---|---|---|
| A269L | 0.039 ± 0.008 | 53 | 4.83 |
| A269F | 0.071 ± 0.004 | 148 | 1.62 |
| G270A | N.A. | 1.01 ± 0.08 | 1.83 ± 0.14 |
| G270S | 1.08 ± 0.27 | 1.00 ± 0.13 | 1.84 ± 0.02 |
| G270F | N.A. | 1.00 ± 0.13 | 1.91 ± 0.09 |
| G270L | N.A. | 1.09 ± 0.05 | 1.83 ± 0.14 |
| L266T | N.A. | 1.33 ± 0.22 | 1.89 ± 0.03 |
| F273A | 0.466 ± 0.074 | 0.336 ± 0.057 | 2.33 ± 0.87 |
| F273W | 3.29 ± 0.76 | 2.60 ± 0.24 | 2.30 ± 0.05 |
| T94I | N.A. | 0.823 ± 0.30 | 3.02 ± 0.03 |
| I189A | 1.51 ± 0.28 | 1.28 ± 0.13 | 1.81 ± 0.02 |
| I189V | 1.17 ± 0.23 | 1.09 ± 0.04 | 1.56 ± 0.02 |
| I189T | N.A. | 0.73 ± 0.18 | 2.35 ± 0.70 |
| I189P | 13.6 ± 2.3 | 9.9 ± 3.7 | 12.9 ± 4.2 |
| Chicken Green | N.A. | 100 ± 8 | 246 ± 17 |
| chGr_P189I | N.A. | 2.7 ± 3.7 | 33± 4.2 |
| I189P_F208W | N.A. | 2.64 ± 0.56 | 16.8 ± 11.7 |
| I189P_F212W | N.A. | 0.60 ± 0.02 | 11.7 ± 7.4 |
| I189P_F212A | N.A. | 27 ± 11 | 42 ± 11 |

**Table S2. The 11CR binding kinetics (×$10^3$ $M^{-1}$ $s^{-1}$) of F208 mutants at different temperatures, measured with Alexa488-based assay**

| | F208A | F208W |
|---|---|---|
| 36 °C | 0.233 ± 0.24 | 0.557 ± 0.33 |
| 32 °C | N.A. | 0.47 ± 014 |
| 28 °C | 0.104 ± 0.30 | 0.333 ± 0.016 |
| 24 °C | N.A. | 0.187 ± 0.010 |
| 20 °C | 0.056± 0.014 | 0.072 ± 0.001 |
| 16 °C | N.A. | 0.081 ± 0.004 |
| 12 °C | 0.014 ± 0.01 | 0.018 ± 0.001 |

**Table S3. The 11CR binding kinetics (×$10^3$ $M^{-1}$ $s^{-1}$) of F212 mutants at different temperatures, measured with Alexa488-based assay**

| | F212A | F212W |
|---|---|---|
| 36 °C | 2.852 ± 0.46 | 0.099 ± 0.012 |
| 32 °C | 2.696 | 0.083 ± 0.003 |
| 28 °C | 1.64 ± 0.25 | 0.064 ± 0.037 |
| 24 °C | 1.270 ± 0.059 | 0.039 ± 0.011 |
| 20 °C | 0.890 ± 0.009 | 0.023 ± 0.010 |
| 16 °C | 0.672 ± 0.39 | 0.019 ± 0.001 |
| 12 °C | 0.319 ± 0.021 | 0.008 ± 0.001 |

**Table S4. The 11CR binding kinetics of F273 mutants ($\times 10^3$ $M^{-1}$ $s^{-1}$) at different temperatures, measured with Alexa488-based assay.**

| | F273A | F273W |
|---|---|---|
| 36 °C | 1.79 ± 0.19 | 10.0 ± 1.5 |
| 32 °C | 0.81 ± 0.24 | 4.33 ± 0.26 |
| 28 °C | 0.466 ± 0.074 | 3.29 ± 0.076 |
| 24 °C | N.A. | 2.12 ± 0.29 |
| 20 °C | 0.333 ± 0.019 | 2.03 ± 0.45 |
| 16 °C | N.A. | 1.314 ± 0.036 |
| 12 °C | 0.113 ± 0.031 | 0.464 ± 0.017 |

**Table S5. The 11CR binding kinetics of I189 mutants (×$10^3$ $M^{-1}$ $s^{-1}$) at different temperatures, measured with Alexa488-based assay**

| | I189A | I189P |
|---|---|---|
| 36 °C | N.A. | 24.8 ± 2.1 |
| 32 °C | N.A. | 14.4 ± 2.5 |
| 28 °C | 1.51 ± 0.28 | 13.6 ± 2.3 |
| 24 °C | 1.038 ± 0.004 | 7.0 ± 0.9 |
| 20 °C | 0.73 ± 0.26 | 4.10 ± 0.16 |
| 16 °C | 0.38 ± 0.13 | 2.69 ± 0.10 |
| 12 °C | 0.235 ± 0.026 | 2.49 ± 0.48 |

**Table S6. The 11CR binding kinetics of I189mutants ($\times 10^3$ $M^{-1}$ $s^{-1}$) at different temperatures, measured with Alexa488-based assay**

| | S144-Alexa488 Rho |
|---|---|
| 36 °C | 2.66 ± 0.44 |
| 28 °C | 1.12 ± 0.25 |
| 20 °C | 0.54 ± 0.12 |
| 12 °C | 0.18 ± 0.4 |
